# Methionine sulfoxide reductase B3 antioxidant activity is indispensable for mouse inner ear cuticular plate structure and hair bundle integrity

**DOI:** 10.1101/2025.03.04.641487

**Authors:** Gowri Nayak, Elodie M. Richard, Byung Cheon Lee, Gavin P. Riordan, Inna A. Belyantseva, Bruno Manta, Thomas B. Friedman, Vadim N. Gladyshev, Saima Riazuddin

**Affiliations:** Department of Developmental Biology, Cincinnati Children’s Hospital Medical Center, Cincinnati, OH, 45229, USA; Department of Otorhinolaryngology Head & Neck Surgery, School of Medicine, University of Maryland, Baltimore, MD, 21201, USA; Division of Genetics, Department of Medicine, Brigham and Women’s Hospital, Harvard Medical School, Boston, MA 02115, USA; Broad Institute, Cambridge, MA, USA; College of Life Sciences and Biotechnology, Korea University, Seoul 02841, Republic of Korea; Laboratory of Molecular Genetics, National Institute on Deafness and Other Communication Disorders, NIH, Bethesda, Maryland, USA; Broad Institute, Cambridge, MA, USA; Department of Molecular Biology and Biochemistry, School of Medicine, University of Maryland, Baltimore, MD, 21201, USA

## Abstract

Methionine sulfoxide reductases (MSR) are enzymes responsible for catalyzing the reduction of methionine-sulfoxides. We previously demonstrated that variants in human *MSRB3*, a member of the MSR family, are associated with profound autosomal recessive prelingual non-syndromic hearing loss DFNB74. To better understand the role of MSRB3 in the auditory pathway, we generated a complete *MsrB3* gene knock-out mouse model of human DFNB74 deafness. The *MsrB3* deficient mouse showed profound hearing loss by postnatal day 16 (P16), which was accompanied by hair cell morphological abnormalities including altered stereocilia bundle shape and cuticular plate degeneration followed by hair cell apoptotic death. Although the absence of MSRB3 primarily affected the actin cytoskeleton, rootlets were present, and the expression and localization of major F-actin stereocilia-core proteins were unaltered. Biochemical assays demonstrated that wild-type MSRB3, but not MSRB3 harboring p.Cys89Gly, the same variant reported for human deafness DFNB74, can repolymerize oxidized actin. Consistent with these *in vitro* data, we observed a decreased ratio of reduced/total actin in the inner ears of *Msrb3* knock-out mice. These data suggest a protective role of MSRB3 in the maintenance and maturation of stereocilia and hair cells, a conserved mechanism aimed at maintaining actin redox dynamics in these sensory cells.

## Introduction

The cochlea is the receptor organ for hearing and consists of rows of sensory and supporting cells that together comprise the organ of Corti. The sensory hair cells perceive sound-induced vibrations through actin-rich structures called the stereociliary bundles on the apical surface of sensory hair cells. Mechanical displacements of the stereociliary bundles are transduced into electrical signals that are then transmitted to the brain. Hair cells, like all other cell types, are susceptible to oxidative stress-induced damage and apoptosis that occur as a result of noise-trauma, ototoxicity or aging^1^. Reactive oxygen species that are released during hair cell activity and metabolism are dealt with by built-in antioxidant mechanisms, failure of which can lead to hearing loss^2–6^.

Surface-exposed methionine residues in proteins are susceptible to reactive oxygen species that can oxidize them to methionine sulfoxide residues resulting in loss or altered biological activity of the protein^7–10^. Methionine sulfoxide reductases are enzymes that catalyze the reduction of methionine sulfoxide to methionine found as both the free-form and incorporated into proteins^9; 11–13^. These enzymes are stereospecific, wherein MSRA is directed to the S diastereomer, methionine-S*-* sulfoxide, while the MSRB family targets methionine-R*-*sulfoxides^14–17^. The cellular distribution of these enzymes varies. In humans, MSRA and MSRB1 localize to the mitochondria and cytosol, MSRB2 to the mitochondria, and MSRB3 isoforms A and B localize to the endoplasmic reticulum and the mitochondria, respectively^18; 19^. Mutations in *MSRB3* were reported to underlie DFNB74 and the protein was shown to localize predominantly to the sensory hair cells in the cochlea^2–4; 6^. *MsrB3* last exon deletion mutant mice and zebrafish morphants had been developed, which exhibited profound hearing loss without vestibular dysfunction, recapitulating the hearing deficit observed in human^6; 20^. In both models, the sensory epithelia show progressive stereociliary degeneration and ultimately hair cell apoptosis. Exogeneous expression of MSRB3, using *in utero* injection of AAV particles into E12.5 mouse embryos, as well as injection of human MSRB3 isoform A mRNA into zebrafish larvae at the one-cell stage, was able to rescue the hearing deficit as well as the stereocilia and hair cells defects^20; 21^. While these models show the pathogenicity of MSRB3 functional deficit and its impact on the auditory pathway, no molecular mechanisms have been proposed to explain the observed phenotype.

The current study investigates the inner ear phenotype of a mutant mouse in which all the coding exons of *MsrB3* were deleted and the functional consequences of the absence of the encoded redox protein in the cochlea. Loss of MSRB3 results in morphological abnormalities in the inner ear hair cells including altered stereocilia bundle shape and cuticular plate degeneration followed by hair cell apoptotic death, that stem from increased oxidative damage to actin molecules in cochlear hair cells. This study highlights the importance of an endogenous redox mechanism in the inner ear for maturation and maintenance of the hair bundle, which is necessary for the mechanotransduction of sound.

## Results

### MSRB3 localizes to hair cells of the organ of Corti

*MsrB3* full gene knock-out mouse was recovered from a cryopreserved strain of KOMP/Velocigene (Figure 1A). To excise the NEO cassette, we crossed the mice acquired from VelociGene with a mouse line expressing the Cre recombinase under the *Zp3* promoter (C57BL/6-Tg(Zp3-cre)93Knw/J)^22^. After Neo cassette removal, loss of MSRB3 expression was confirmed by quantitative PCR and immunolabeling (Figure 1B-C). Capitalizing on the insertion of a LacZ reporter cassette downstream of *MsrB3* promoter, we assessed the spatial and temporal expression pattern of *MsrB3* in the inner ear. *MsrB3* expression was noted specifically in hair cells of the cochlear sensory epithelium from early postnatal time points until two months of age, the oldest time point studied here (Figure 1D-E). Reporter expression was also observed in the spiral ganglion and the spiral ligament (Figure 1D).

**Figure 1.**
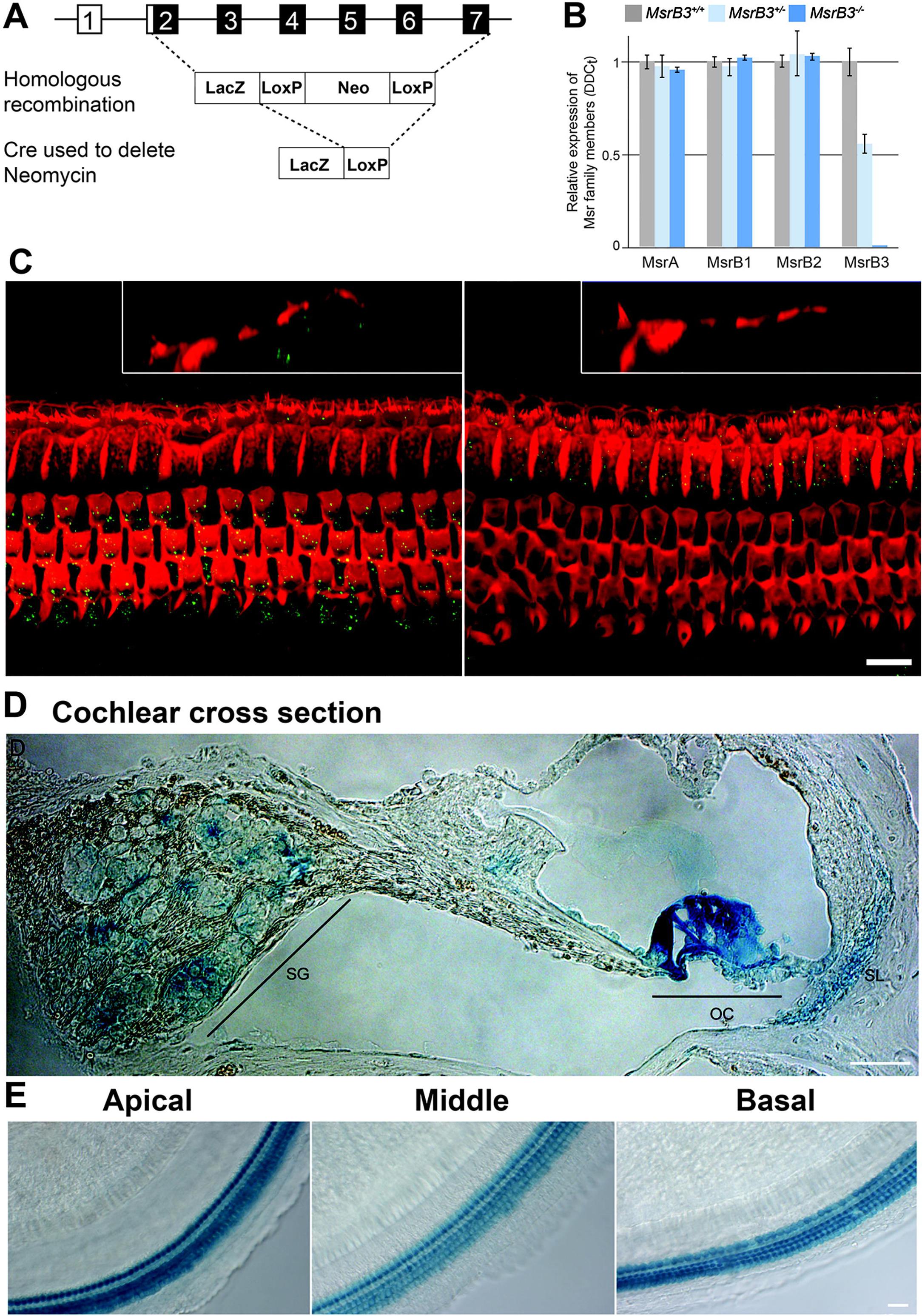
Generation and validation of *MsrB3* knockout mice. (A) Schematic representation of generation of *MsrB3^-/-^* mice. Exons 2 to 7 of *MsrB3* were replaced by homologous recombination with a LacZ and Neomycin cassette. The Neomycin resistance cassette (Neo), flanked by LoxP sites, was removed by crossing the mice with mice expressing a Cre recombinase gene (C57BL/6-Tg(Zp3-cre93Knw/J; zona pellucida 3 gene). (B) Quantification of *MSR* family members expression, normalized against *Gapdh*, in the cochleae of *MsrB3^-/-^* mutant mice and their control littermates (*MsrB3^+/+^* and *MsrB3^+/-^*) at P0. As anticipated, *MsrB3* expression is reduced in heterozygous mice and undetectable in homozygous mutant mice. No variation of expression was observed in any other genes of the *MSR* family (means ± SEM). (C) Maximum intensity projections of confocal Z-stacks of whole-mount cochleae at P16, labeled with MSRB3 antibodies (green) and phalloidin (red) are shown. Insets in both images show a y-z projection of the organ of Corti’s epithelium. While in the wild-type mouse MSRB3 protein (green) is localized in the cytoplasm of the outer and inner hair cells, no immunolabeling was detected in the *MsrB3^-/-^* mutant mice (C, right panel). Scale bar: 10µm. (D-E) X-gal staining was performed on *MsrB3^+/-^* mice to document *MsrB3* promoter activity in the organ of Corti from P6 mice. *MsrB3* is expressed in hair cells throughout the cochlear turns along with the spiral ganglion (SG) and spiral ligament (SL) regions. Scale bars: 15 µm.

### *MsrB3* knock-out mouse has an early onset profound deafness

The cochlear function of *MsrB3^-/-^* and control (*MsrB3^+/+^*, *MsrB3^+/-^*) mice was evaluated by measuring auditory brainstem responses (ABR) using broadband click and tone-burst sounds at P16. While *MsrB3^+/+^*and *MsrB3^+/-^* mice had normal and indistinguishable ABR thresholds, *MsrB3^-/-^* mice did not respond to click or tone burst stimuli of even a 100 dB sound pressure level (SPL), indicating that they are profoundly deaf (Figure 2A). Distortion product otoacoustic emissions (DPOAEs), which represents the outer hair cell (OHC) function, was found to be reduced to no responses in the mutant mice, although the responses were often not discernible from the noise floor (Figure 2B). The ABR and DPOAE assessments suggest that the absence of MSRB3 leads to cochlear deficiencies that result in hearing loss. Next, mechanotransduction of cochlear hair cells was evaluated at P5 for control and *MsrB3^-/-^* mutant mice. We observed no differences in the uptake of the channel-permeable fluorescent styryl dye FM1-43 (Figure 2C), suggesting that *MsrB3* deficiency is unlikely to cause mechanotransduction defects.

**Figure 2.**
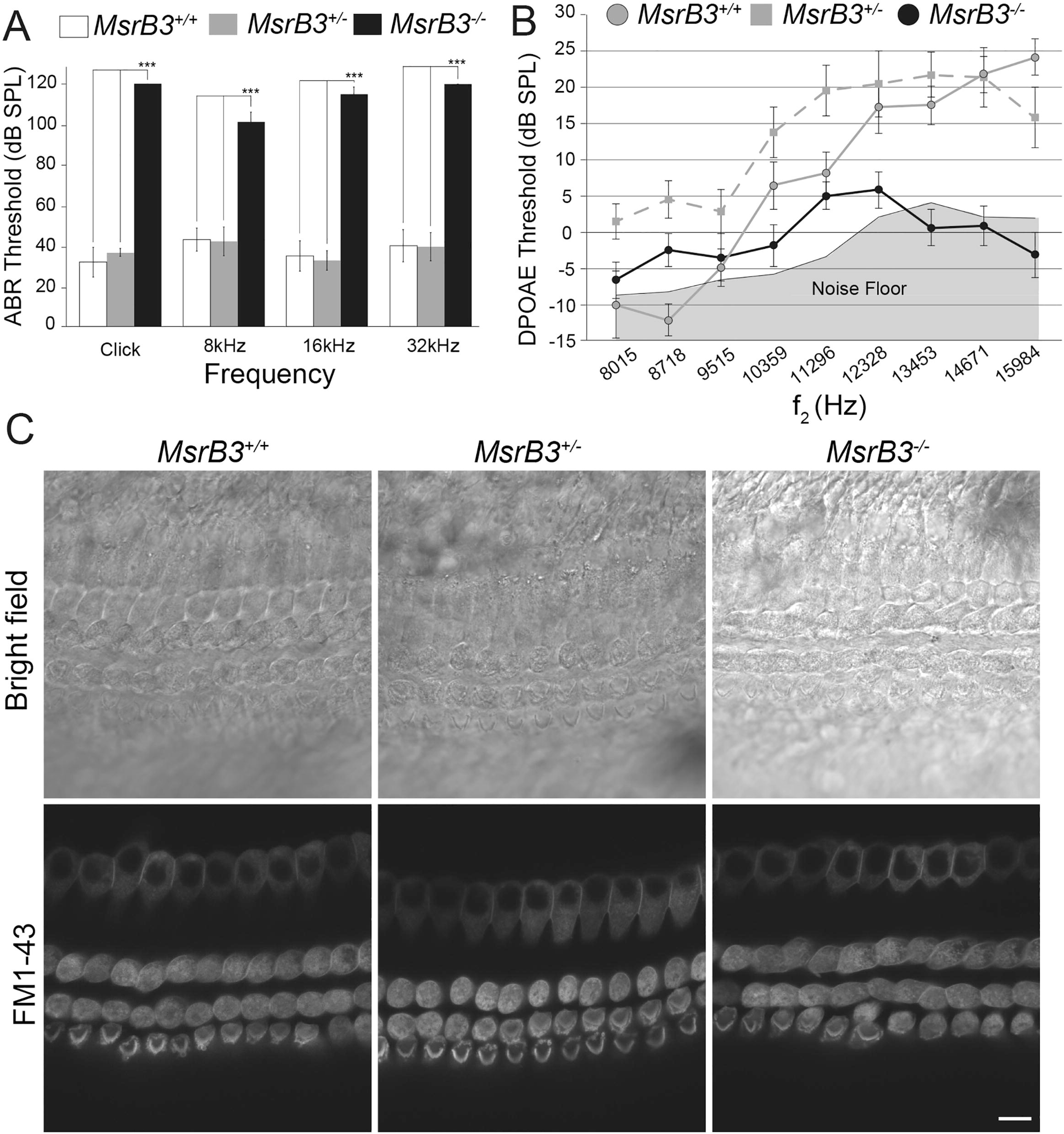
*MsrB3* mutant mice are profoundly deaf by P16 with likely intact MET channel function. (A) Averaged auditory brainstem responses (ABR) thresholds of *MsrB3^-/-^*mice (n= 9) and their control littermates (*MsrB3^+/-^* (n=9) and *MsrB3^+/+^* (n=7)) at P16 in response to broadband click stimuli and 8, 16 and 32kHz tone-bursts, with sound pressure level (SPL) of 0 to 120 dB. The *MsrB3^-/-^* mice are profoundly deaf and showed statistically significant (***p<0.001) elevated thresholds for all the tested frequencies compared to their control littermates (mean ± SEM). (B) Distortion product otoacoustic emissions (DPOAE) thresholds of *MsrB3^-/-^* (black circle, n=9), *MsrB3^+/-^* (gray square, n=9) and *MsrB3^+/+^*(grey circle, n=7) at P16, represented as a function of *f2* stimulus frequencies. *MsrB3^-/-^* mice showed reduced to no responses, with values close to the noise floor, suggesting that their OHC at this age are non-functional (mean ± SEM). (C) Mechanotransduction was assessed by FM1-43 dye uptake on *MsrB3^-/-^* and control littermates mice explants at P5. No apparent difference in the uptake of the FM1-43 dye was observed between the mutant and the control mice. Scale bar: 5 µm.

### Loss of MSRB3 results in early postnatal developmental defects in the organ of Corti

To determine the root cause of hearing deficiency, we evaluated the gross morphology and cytoarchitecture of the organ of Corti in *MsrB3^-/-^* mutant mice. The cochlear sensory epithelium in the mutant animals appeared normal at birth (Supplementary figure 1). However, detectable abnormalities were seen by P4, including curving of the outer edges of the outer hair cell (OHC) stereociliary bundle, indicating that positions of peripheral stereocilia within a bundle and their rootlets are deviated from the normal positions within the cuticular plate likely due to the altered rootlet to actin meshwork connections or actin meshwork structure of the hair cell cuticular plate (Figure 3). These OHC changes were mostly restricted to the middle and basal turns of the cochlea, while all inner hair cells (IHCs) and all apical turn OHCs looked normal, consistent with near normal DPOAE data at 8-12 kHz region (Figure 2B). By P10, however, phenotypic changes were seen throughout the cochlea when visualized with a β-spectrin antibody, which in control mice uniformly labelled the cuticular plate except for the region of the kinocilium (Figure 3). In nearly all mutant OHCs, however, several “β-spectrin-free” zones near the cuticular plate edges were noted, which likely indicated a disrupted cuticular plate structure to which stereocilia rootlets are anchored^23^ (Figure 3). While some hair cell death was obvious at this stage, the cochlear sensory cells were largely preserved until approximately P16 (Figure 3). By three weeks of age, the basal-most end of the cochlea was almost devoid of OHCs, while IHCs were still present. The organ of Corti in the remaining turns of the cochlea also frequently had missing or apoptotic hair cells at this stage (Figures 3 and 4A).

**Figure 3.**
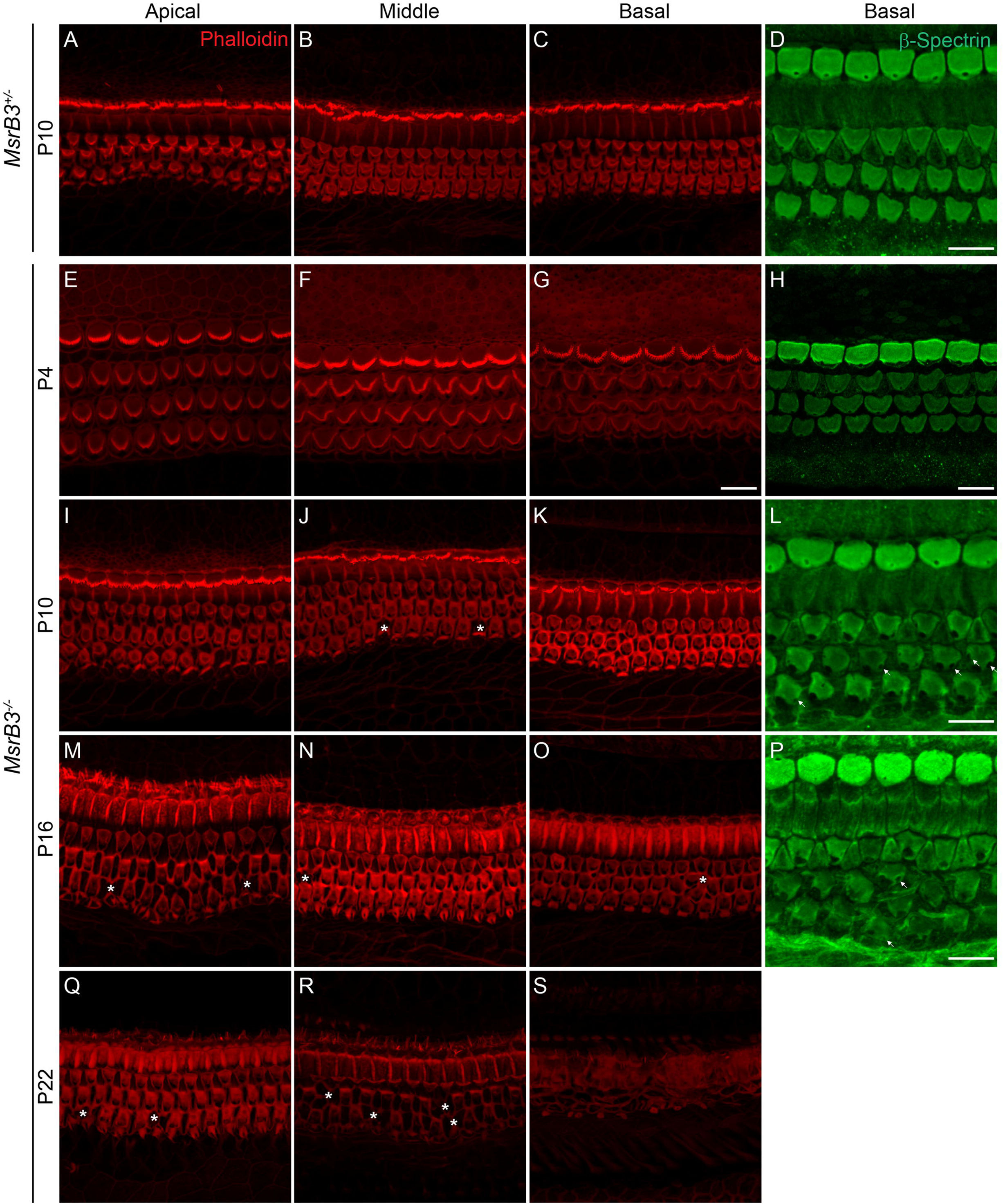
Organ of Corti’s hair cells of *MsrB3* mutant mice showed abnormal stereociliary bundles, cuticular plate defects, and progressive hair cells degeneration. Maximum intensity projections of confocal Z-stacks of whole-mount cochleae labeled with anti-β-spectrin (green) and phalloidin (red) are shown. (A-C) Representative images from the apical, middle and basal turns of the organ of Corti of a control mouse (*MsrB3^+/-^*) at P10. (D) Representative image of hair cells cuticular plates labeled with β-spectrin from the basal turn of the organ of Corti of a control mouse (*MsrB3^+/-^*) at P10. (E-S) representative images of the organ of Corti from the three turns of the cochleae of *MsrB3^-/-^*mice at P4 (E-H), P10 (I-L), P16 (M-P), and P22 (Q-S). Panels H, L, and P show representative images of hair cells cuticular plates labeled with β-spectrin from the basal turn of the organ of Corti of mutant mouse (*MsrB3^-/-^*) at P4, P10, and P16 respectively. Compared to control mouse, *MsrB3^-/-^* mice showed abnormal stereocilia bundles, starting at P4, in middle and basal turns (F-G). By P10, some signs of hair cell loss are visible (asterisks, J) associated with abnormal holes (arrows) in the spectrin labeled cuticular plate (L). The hair cell loss worsened rapidly and by P22, all hair cells of the basal turn were degenerated, and both middle and apical turns showed some hair cell loss (asterisks). Scale bars: 10 µm. Scale bar in (S) applies to all images except otherwise mentioned. Scale bar in (G) applies for (E) and (F). (*) indicates missing hair cell.

**Figure 4.**
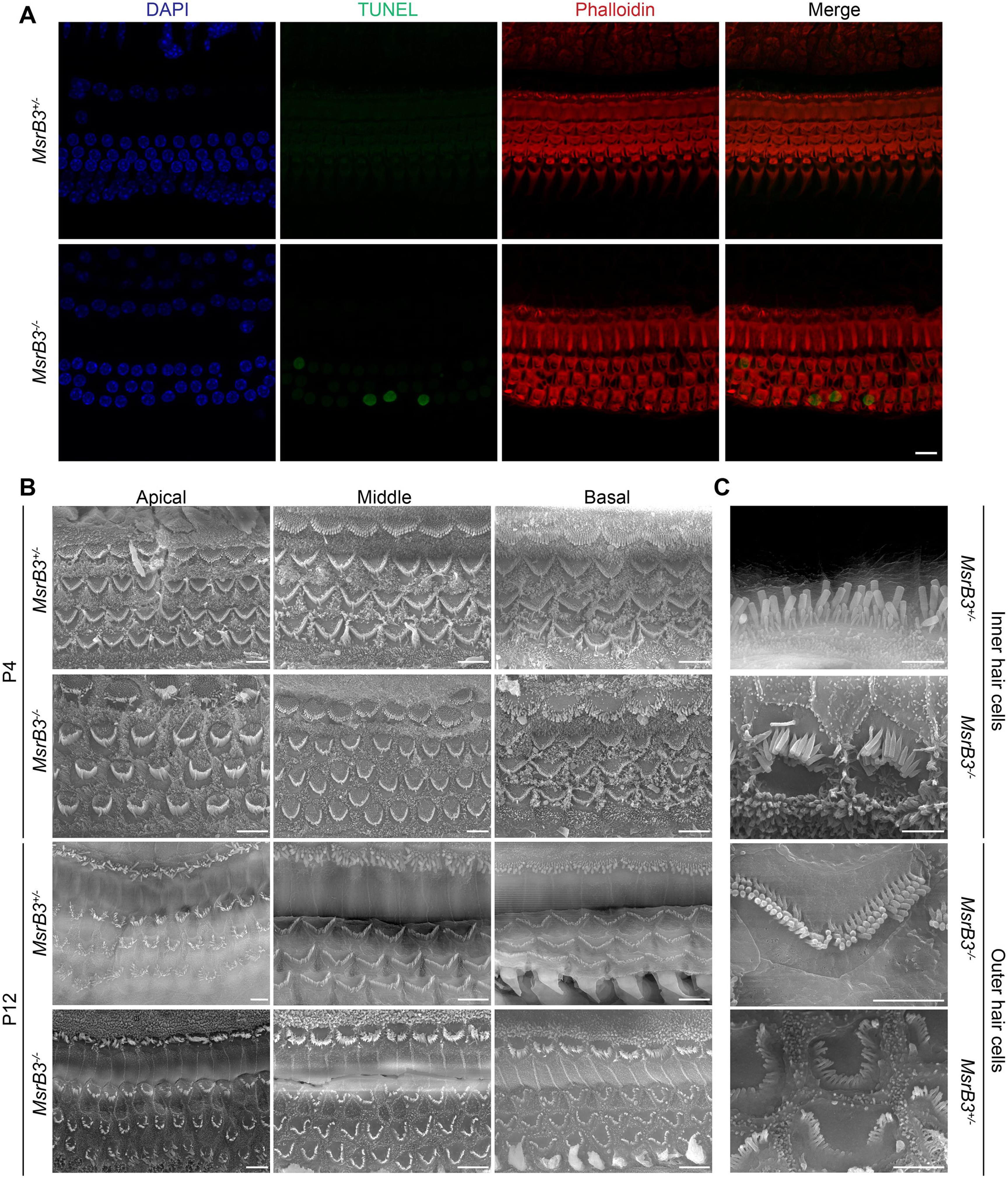
Loss of MSRB3 induces apoptosis and disrupts the stereocilia bundles morphology. (A) TUNEL assay (green) was performed on whole-mount cochleae of *MsrB3^+/-^* control and *MsrB3^-/-^* mice at P20. Nucleus were highlighted by DAPI (blue) and actin filaments were stained using Phalloidin (red). Only hair cells in *MsrB3^-/-^* mutant mice were TUNEL positive. Scale bar: 5µm and applies to all panels. (B) Scanning electron micrographs of the organ of Corti from the three turns of the cochleae of *MsrB3^-/-^* and the heterozygous control mice at P4 and P12. Starting in the basal turn, the stereocilia bundles of the *MsrB3^-/-^* mice lost the characteristic “V” shape seen in the controls to an “omega” shape, as early as P4 and progresses rapidly. By P12, most of the stereocilia bundles in the middle and basal turns of the cochlea exhibit an “omega” shape. Scale bars: 5 µm. (C) High magnification images of representative inner and outer hair cells stereocilia bundle morphology of *MsrB3^-/-^* and control mice at P12. Scale bars: 2 µm.

Next, we evaluated the changes in the shape of the apical surface of hair cells and stereocilia bundles by scanning electron microscopy (SEM). At P4, especially in the basal turn of the cochlea, the stereocilia bundles of OHCs in the *MsrB3^-/-^* mice lost the characteristic “V” shape (Figure 4B). These changes progressed rapidly and by P12, most of the stereocilia bundles in the middle and basal turns of the cochlea exhibited an “omega” shape (Figure 4B-C). The stereocilia were no longer upright but angled away from the basal body region (Figure 4B-C). In some OHCs, the stereocilia were in various stages of dissolution and the apical surfaces of most hair cells had a more rounded appearance compared to those of control cells (Figure 4B-C). The curving of the stereociliary bundles and the subsequent progressive collapse of the stereocilia suggested an anchoring defect of the actin filaments or overall destabilized actin structure. Transmission electron microscopy (TEM) imaging revealed seemingly intact rootlets, without any obvious structural alterations in P14-P21 mutant mice (Supplementary figure 2), although this observation cannot rule out a potential anchoring deficits due to loss of MSRB3 function. Occasionally, we observed some irregularities in the P14-P21 outer hair cell cuticular plate density (Supplementary figure 2). Thus, the stereocilia structural abnormalities noted in *MsrB^-/-^* mice could be due to structural abnormalities in the cuticular plate actin meshwork. However, despite widespread defects in hair bundle shape, the localization of several apically expressed proteins including Ezrin, Radixin, TRIOBP, and T Plastin^24^, remained unaltered in the postnatal cochlea (Figure 5).

**Figure 5.**
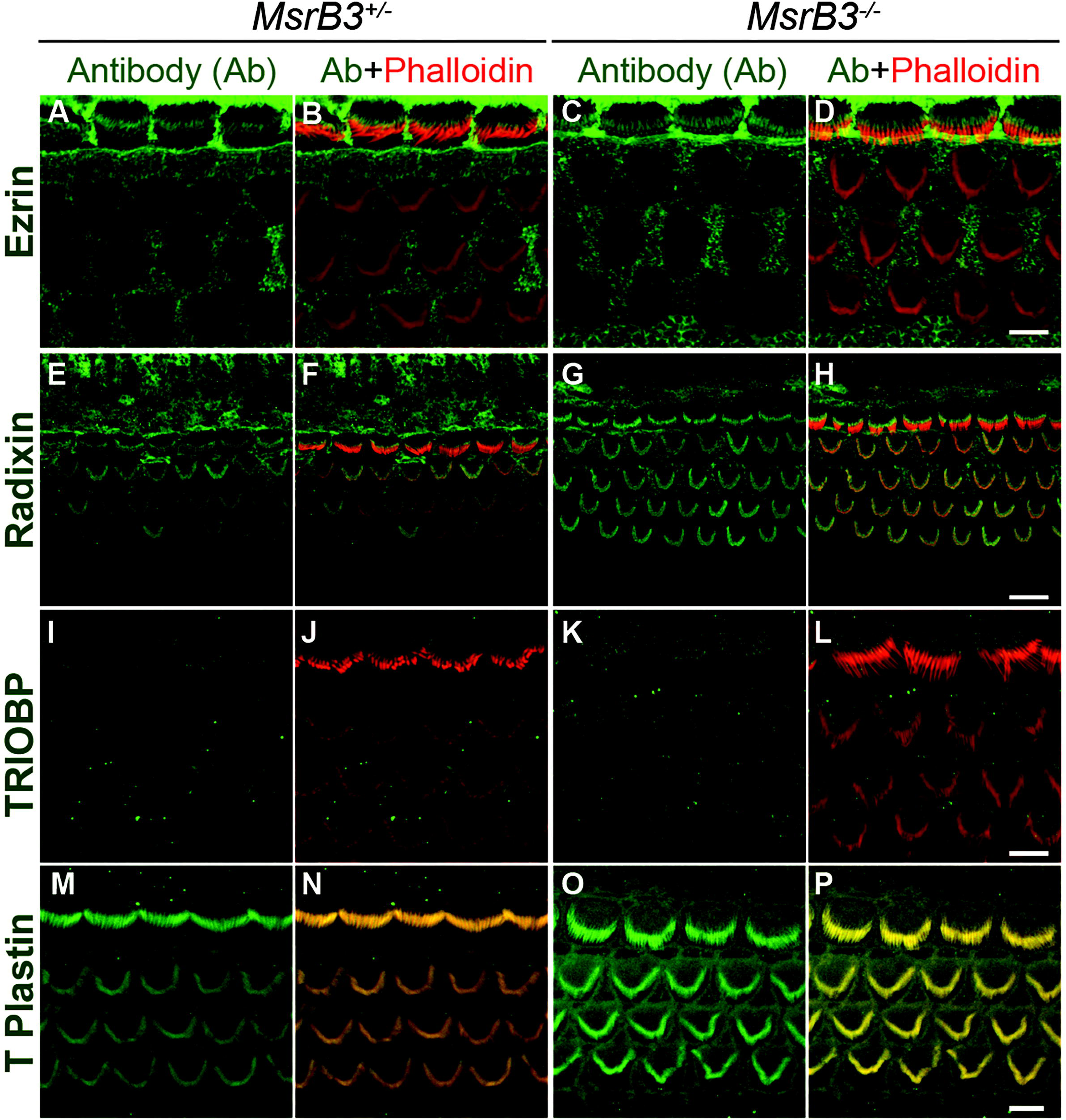
Actin-core related proteins do not show an apparent change in their localization in the hair cells of *MsrB3^-/-^* mutant mice. Maximum intensity projections of confocal Z-stacks from the middle turn of the organ of Corti whole-mount cochleae labeled with antibodies to Ezrin (A-D, green), Radixin (E-H, green), TRIOBP4/5 (I-L, green), T-plastin (M-P, green) and counterstained by rhodamine-phalloidin (red) are shown for heterozygous and *MsrB3* mutant mice at P7. Scale bars shown in (D), (L) and (P) are 5 µm and apply to panels (A, B, and C), (I,J, and K) and (M,N, and O), respectively. Scale bar in (H) is 10 µm and applies to panels (E), (F), and (G).

### Catalytically active MSRB3 can re-polymerize actin *in vitro*

A prior study reported the role of MSRB1, another member of the MSR family, in facilitating actin assembly, specifically by reducing the oxidized methionine in actin, brought about by the monooxygenase MICAL^25^. Thus, the reversible oxidation of actin by MICAL prevents filamentous actin assembly, which is antagonized by MSRB1. To determine if MSRB3 is also able to stereo-selectively targeting oxidized methionine in actin, the repolymerization of MICAL-treated, pyrene-labelled actin was tested in the presence of full-length wild-type human MSRB3 (MSRB3^WT^) or a variant containing the p.Cys89Gly substitution (MSRB3^MUT^) that underlies human deafness DNFB74^2^. While MSRB3^WT^ was able to repolymerize oxidized actin as measured by the increase in fluorescence intensity, loss of the catalytic p.Cys89 abolished this activity (Figure 6A). This observation strongly suggests that the hair bundle and cuticular plate phenotypes seen in the mutant mice are likely due to an accumulation of G-actin molecules with oxidized methionines, which are incapable of re-polymerizing, an essential feature of hair bundle function and maintenance.

**Figure 6.**
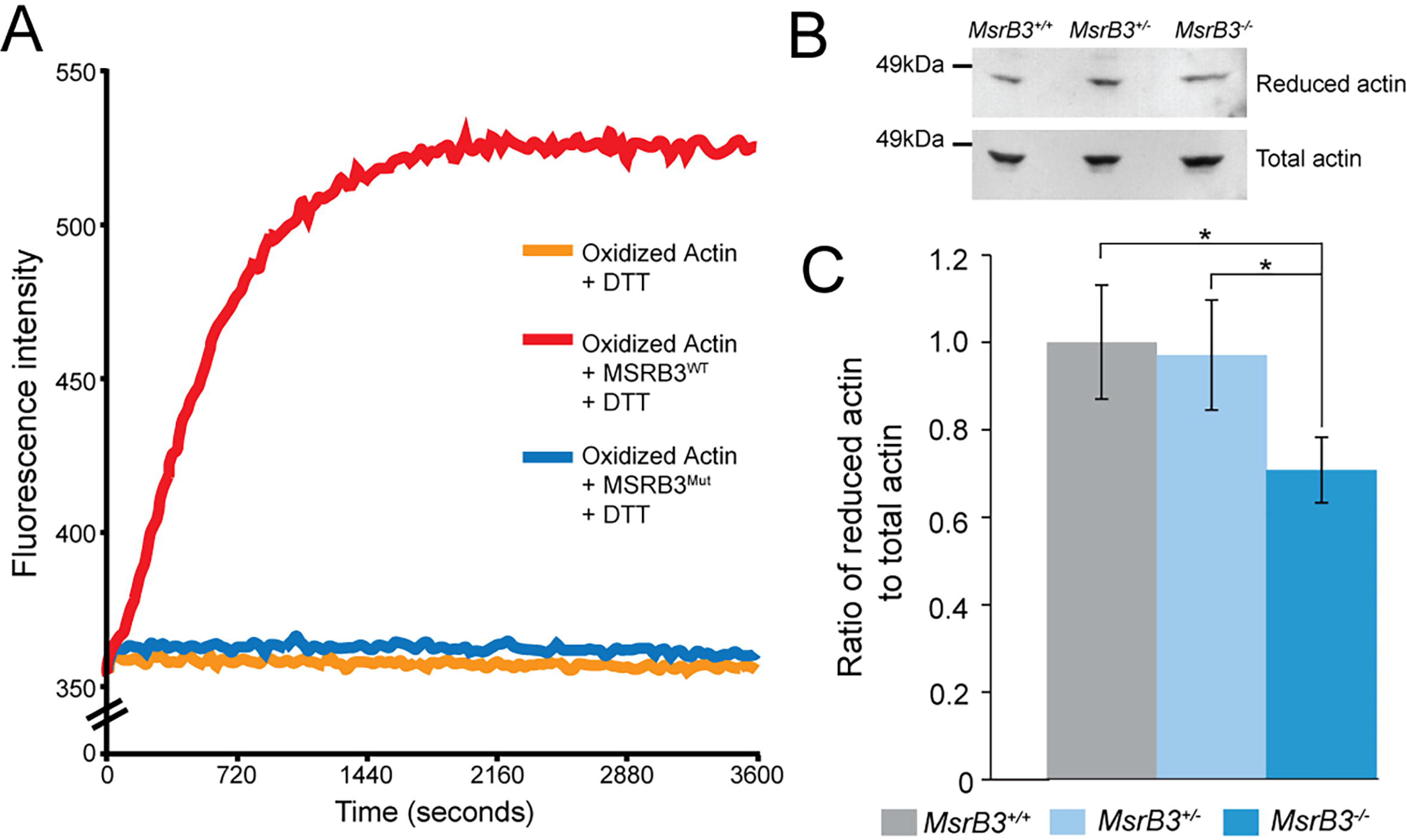
MSRB3 promotes actin polymerization and the absence of MSRB3 in mutant mice affects the reduced/oxidized actin ratio in the inner ear. (A) Pyrene-labeled actin monomer was monitored for polymerization by assaying changes in fluorescence at 407 nm (excitation at 365 nm) in the presence of wild type human MSRB3 (red, MSRB3^WT^) or deafness associated protein MSRB3 p.Cys89Gly (blue, MSRB3^Mut^). As a control, fluorescence intensity of the preassembled pyrene-labeled actin was monitored in the absence of MSRB3 (orange). (B) Representative western blot showing the amount of reduced and total actin in the whole inner ear tissue lysates from *MsrB3^-/-^* and control littermates (*MsrB3^+/-^* and *MsrB3^+/+^*) at P14, using antibodies specific for the reduced form of actin and total actin. (C) The ratio of reduced to total actin was calculated by using ImageJ to quantify the band densities shown (B). All data were normalized to the *MsrB3^+/+^* ratio (means ± SEM). This experiment was repeated independently four times with cochleae from three mice per condition. *MsrB3^-/-^* mutant mice showed a statistically significant (Student’s *t* test, * p<0.05) decreased ratio of reduced to total actin suggesting a possible role for MSRB3 in the regulation of reduced actin to maintain the stereocilia structure in the inner ear.

### *MsrB3* mutant mice have a lower level of reduced actin

The hypothesis that the *MsrB3^-/-^* mutant mice accumulate G-actin with oxidized methionines was tested by quantifying the ratio of the reduced form of actin to total actin. Using antibodies raised against reduced actin and total actin, the levels of the respective actin pools in the inner ear tissue samples of control and *MsrB3^-/-^* mutant mice were determined by western blotting. The amount of reduced actin in the normal hearing *MsrB3^+/-^* inner ears of was found to be like that seen in wild-type animals. However, *MsrB3^-/-^*inner ears had significantly lower levels of reduced actin (Figure 6B-C). Thus, we posit that the hair bundle defects seen in the *MsrB3* mutant mice are likely to be a result of elevated oxidative damage that targets the actin, and likely other proteins, and affects their dynamics and function in the cochlear sensory cells.

## Discussion

MSRs are enzymes that repair damaged proteins by selectively reducing oxidized methionine residues^26; 27^. Among the four paralogs (MSRA, MSRB1, MSRB2, MSRB3), so far, only MSRA and MSRB3 have been associated with hearing loss^2; 5^. While *MsrA* knock-out mice present a progressive hearing loss, starting at the highest frequencies^5^, *MSRB3* variants in humans lead to pre-lingual bilateral profound hearing loss DFNB74^2–4^. As has been observed before in *MsrB3* exon 7 mutant mice^20^, our analysis of the cochlear phenotype of a *MsrB3* null mouse line also revealed hair bundle abnormalities: a characteristic “omega” shape of the stereocilia bundle as early as P4 in the basal turn of the organ of Corti similar to the stereocilia shape observed in Triobp-5 and ANKRD24 knockout mice^28; 29^. In addition, *MsrB3* knockout mice exhibited progressive hair cell loss, starting with OHCs at the basal turn of the cochlear coil, until the sensory epithelium was completely degenerated by P60 (data not shown). TUNEL immunoassays revealed that the hair cell loss observed in the mutant mice was due to apoptotic cell death (Figure 4A).

As the first sign of deficit observed in the mutant hair cells is the collapse of the stereociliary bundle, we first focused on the three main components of the stereocilia that ensure its stability: the actin-core related proteins, the rootlets and the cuticular plate. Stereocilia are composed of an F-actin core that contains crosslinked, parallel actin microfilaments and actin cytoskeleton-associated proteins that have proven crucial for development, maintenance and stabilization of the stereocilia^30^. However, we observed no obvious differences in the expression of several of the actin-core associated proteins in mutant mice (Figure 5).

At the tapered end of the stereocilium, the actin filaments form a rootlet that anchors a stereocilium in the actin-rich meshwork of the cuticular plate^31; 32^. Rootlets provide rigidity as well as flexibility to stereocilia, which are essential for optimal sensitivity of the hair bundle ^31; 32^. Genetic studies have recently shed light on the central role of TRIOBP, an F-actin-bundling protein, in rootlet formation^28; 33^. Interestingly, *Triobp* mutants exhibit a phenotype similar to *MsrB3* knock-out mice, where the hair bundle develops normally at first but soon after the onset of hearing, the stereocilia fuse together, and the hair cell degenerate rapidly. In the *Triobp* isoform 4/5 deficient mouse models, the phenotype is explained by the absence of rootlets, which makes the stereociliary bundles fragile^28; 33^. However, *MsrB3* deficient mice did not exhibit any rootlet deficit at P14-P21 (Supplementary figure 2), well beyond the age at which the hair bundle phenotypes are first seen. Finally, although the “potholed” appearance of the cuticular plate is secondary to observed hair bundle damage, it is apparent before the onset of hearing and is likely to be a leading cause of hearing loss in these mice.

We then questioned if MSRB3 was involved in the maintenance and/or maturation of the stereocilia bundles. Indeed, hair cells of *MsrB3* deficient mice seem to develop normally, initially, without any morphological aberration (Supplementary Figure 1) and exhibit normal uptake of the styryl dye, FM1-43, indicative of a functional mechanotransduction apparatus (Figure 2C). However, morphological abnormalities in the stereocilia and hair cell degeneration are seen within the first two weeks of life, which accelerate with the maturation of the sensory epithelium and acquisition of hearing (Figures 3 and 4). These results suggest possible developmental abnormalities of stereocilia and/or the cuticular plate in which stereocilia are anchored in mutant mice. Our data show that MSRB3 is necessary for the structural integrity of the cuticular plate essential for normal stereocilia anchoring and maintenance of the stereociliary bundle.

The regulation of actin dynamics, through rapid turnover of depolymerized/polymerized actin is essential for maintenance of stereocilia. While two contradictory processes have been proposed: a continuous depolymerization of the actin filaments at the base with re-polymerization at the tip^34^, and continuous turnover at the tip of the stereocilia exclusively^35–37^, both mechanisms agree on the essential role of actin dynamics in the maintenance of the stereocilia. The transition between globular (G) and filamentous (F) actin and its dynamics are regulated by many actin-associated proteins. Among others, the role of MICAL, a monooxygenase has been highlighted previously. In Drosophila, two Met residues in actin (46^th^ and 49^th^ methionines) can be selectively oxidized by MICALs, inhibiting F-actin assembly^38; 39^. In humans, MICAL homologs were also found to regulate actin microfilaments and actin stress fibers^40^, strengthening the implication of MICAL-dependent oxidation in regulation of actin dynamics. In 2013, Lee et. al. linked the activity of MICAL protein to two of the MSRB proteins: MSRB1 and MSRB2^25^. Contrary to MSRA, these proteins can re-polymerize G-actin that is previously disassembled by MICAL. These MSRB proteins work in a stereospecific manner with MICAL, regulating the actin assembly turnover. We hypothesized that MSRB3 may have the same MICAL antagonist effect and promote the reassembly of oxidized actin through its reduction. Using the test standardized for MSRB1 and MSRB2 previously^25^, we demonstrated that MSRB3 protein is indeed able to repolymerize oxidized actin (Figure 6A), indicating a role for MSRB3 in the regulation of actin dynamics in the hair cell cuticular plate, maintaining its integrity. Interestingly, MSRB3 harboring the most reported variant (p.Cys89Gly) associated with human deafness^4^ DFNB74 is unable to re-polymerize G-actin.

While other MSR proteins have been largely involved in oxidative stress protection in mammals^41–43^, the protective role of MSRB3 in redox homeostasis is yet to be determined. We did not observe signs of major oxidative stress in the inner ear of the mutant mice compared to their control littermates (Supplementary figure 3), as also previously reported by others^6^. Importantly, none of the other MSR family genes were dysregulated in the cochlea of P0 *MsrB3^-/-^* (Figure 1B), suggesting a unique role of MSRB3 that cannot be compensated by MSR paralogs. Based on the MSRB3/MICAL antagonist effect on actin dynamics, we speculated that MSRB3 might have a protective role on actin molecules to reverse oxidative damages. In support of this, we found that the inner ears of knockout mice had a decreased ratio of reduced actin to total actin compared to that seen in wild-type mice, suggesting that among all MSR proteins, MSRB3 is particularly critical for reversing oxidation of actin molecules in the inner ear. Taken together, these data support the role of MSRB3 in actin polymerization and dynamics, through its redox status. In the absence of MSRB3, there is less repolymerization of oxidized actin occurs leading to the disruption of the cuticular plate actin meshwork as well as altered stereocilia positions at the edges of a bundle leading to an “omega” shape of the stereociliary bundle. In addition, using traditional TEM it is not practical to observe entire length and width of all rootlets of a hair cell stereocilia simultaneously and we cannot rule out some rootlet abnormalities that could be revealed by a follow-up study of rootlet development in MSRB3 mutant mice using FIB-SEM with 3D-reconstructions. In conclusion, this study establishes a specific role of MSRB3 in the inner ear. MSRB3 maintains hair cell through its redox mechanism to preserve normal actin dynamics in the cuticular plates of the cochlear sensory cells, which is prerequisite for the normal function of stereocilia bundles.

## Methods

### Mutant mice and genotyping

The *MsrB3* knock-out mice were recovered from a cryopreserved strain from KOMP/Velocigene (http://www.velocigene.com/komp/detail/12720). In this model, coding exons 2 to 7 were replaced by a LacZ and NEO cassettes. To excise the NEO cassette, and avoid any potential phenotype linked to its presence, we crossed the mice acquired from Velocigene with a mouse line expressing the Cre recombinase under the *Zp3* promoter (C57BL/6-Tg(Zp3-cre)93Knw/J, Jackson Laboratories, #003651). The females, from F1 generation, were then crossed with wild-type male (C57BL/6J) and the F2 generation was screened for excised alleles in both sexes.

### ABR and DPOAE

Hearing function was evaluated by ABR analyses at P16 of *MsrB3^+/+^* (n=7), *MsrB3^-/+^* (n=9), and *MsrB3^-/-^* (n=9) male mice. Mice were anesthetized with intraperitoneal injections of 2.5% Avertin (0.015 ml/g body weight). All recordings were done in a sound-attenuated chamber using an auditory evoked potential diagnostic system (Intelligent Hearing Systems, Miami, FL) with high frequency transducers, as previously described^44^. Responses to 50 µs duration clicks, and 8, 16 and 32 kHz tone bursts were recorded. Thresholds were determined in 5- or 10-dB steps of decreasing stimulus intensity, until waveforms lost reproducible morphology. The maximum sound intensity tested for each frequency was 110 dB SPL.

DPOAEs were recorded from *MsrB3^+/+^* (n=7), *MsrB3^-/+^* (n=9), and *MsrB3^-/-^* (n=9) male mice at P16 with an acoustic probe (ER-10C, Etymotic Research) using DP2000 DPOAE measurement system version 3.0 (Starkey Laboratory). Two primary tones, with a frequency ratio of *f*_2_/*f*_1_ = 1.2, where *f*_1_ represents the first tone and *f*_2_ represents the second, were presented at intensity levels *L*_1_ = 65 dB SPL and *L*_2_ = 55 dB SPL. *f*_2_ was varied in one-eighth octave steps from 8 to 16 kHz. DP grams comprised 2*f*_1_–*f*_2_ DPOAE amplitudes as a function of *f*_2_.

### Immunofluorescence

Inner ears were fixed with 4% paraformaldehyde, overnight at 4°C and permeabilized with pre-block using 0.2% Triton-X100 in 10% heat-inactivated horse serum for 1 hour. The paraformaldehyde-fixed cochleae were probed overnight with primary antibody and after three washes, were probed with the secondary antibody for 1 hour. For all hair cell body staining, rhodamine-conjugated phalloidin (Invitrogen) was used at 1:300 dilution to visualize the cuticular plate and hair bundles. Primary antibodies against MYO7a (Proteus) were used at 1 in 250, β-spectrin (BD Biosciences) at 1 in 250, MSRB3 antibody (Sigma) at 1 in 50, Ezrin (Abcam) at 1 in 50, Radixin (Abcam) at 1 in 50, TRIOBP4/5 (Friedman lab^33^) at 1 in 50 dilutions. Alexa 488 Goat anti-rabbit and Alexa 488 Goat anti-mouse secondary antibodies (Invitrogen) were used at 1 in 500 dilution. When necessary, DAPI was used at 20 µg/ml. After every secondary antibody incubation, the tissues were washed thrice in PBS and mounted on slides using Vectashield ® mounting medium (Vectorlabs) and viewed under a LSM meta 510 confocal microscope using a 63x, 1.3 N.A. oil-immersion objective.

### Scanning electron microscopy analysis

Inner ears were removed from animals after decapitation, and the cochleae were exposed. A small piece of cochlear capsule was removed from the cochlear apex and the entire bulla was immersed in fixative (2.5% glutaraldehyde, 0.1 M sodium cacodylate containing 2 mM CaCl_2_ for 1.5 hours). After three quick washes with 0.1 M sodium cacodylate buffer, the inner ears were post-fixed in 1% osmium tetroxide in 0.1M sodium cacodylate buffer for 1 hour at room temperature. The inner ears were washed thrice with phosphate buffered saline (PBS) buffer before leaving in PBS containing 0.25 M EDTA for 2 days at 4°C. The samples were then microdissected in water to remove the stria vascularis and the spiral ligament was cut-off to expose the organ of Corti. The cochlear tissues were then dehydrated using ethanol gradient, critical-point dried, sputter-coated with gold and imaged on a scanning electron microscope (Hitachi SU3900). Where necessary, the images were adjusted for optimal brightness and contrast using Photoshop® creative studio CS5.1 (Adobe).

### Transmission Electron Microscopy

Inner ears of P14 and P21 *MsrB3* homozygous wild-type and mutant mice were dissected and fixed using 2.5% glutaraldehyde and 2% paraformaldehyde (Electron Microscopy Sciences) in 0.1M phosphate buffer for 2 hours at room temperature, washed in 1X phosphate buffer, decalcified in 100 mM EDTA in 1X phosphate buffer for 48-72 hours at 4°C, washed in phosphate buffer, dehydrated in increasing concentrations of ethanol, poststained in 1% osmium tetroxide and embedded into epon-812 resin (Electron Microscopy Sciences) as previously described (Katsuno et al., 2019). Ultrathin sections ∼70-90 nm were cut using ultracut UCT-7 (Leica), post-stained with 1% Uranyl acetate (Electron Microscopy Sciences) and lead citrate (Sigma-Aldrich) and visualized using JEOL1200 transmission electron microscope (JEOL).

### Western blots

Inner ears were dissected from two mice for each of the indicated genotypes at P14. The tissues were homogenized in 200 µl of lysis buffer [0.05M Tris pH 7.5, 150mM NaCl, 0.1% NP40 and cocktail protease inhibitor (Sigma)] and sonicated to sheer the DNA followed by centrifugation at 14000 rpm for 20 minutes at 4°C to pellet the cellular debris. The supernatants were collected in fresh tubes and ¼ volume of 4x Laemmli sample buffer without any reducing agent were added. The samples were heated for 12 minutes at 70°C and centrifuged before resolving 25 ul of each sample on a 10% SDS-PAGE gel. After electrophoresis, the samples were electrophoretically transferred onto nitrocellulose membranes by the semi-dry Western blotting method. The blots were pre-blocked in 5% low-fat dried milk powder in Tris-buffered saline containing 0.05% Tween-20 before incubating with pan actin antibodies (Cytoskeleton Inc, #AAN01) at 1 in 1000 and reduced actin antibodies (described previously^25^) at 1 in 1000. After three washes, the blots were incubated with horse-radish peroxidase conjugated goat anti-rabbit secondary antibody (GE Healthcare) and the signals were detected using the Enhanced Western blotting kit (Piercenet). Alongside all the samples, a Western molecular standard marker was used to determine approximate molecular weights.

### Semi-quantitative expression studies (RT-PCR)

RNAs from inner ear were extracted at P0 from *MsrB3* wild-type, heterogygous and homozygous mice using the Ribopure kit (Ambion). SMARTScribe Reverse Transcriptase kit (Clontech) was used to generate cDNAs, and SYBRgreen technology (Qiagen) was used to perform the qPCR using genes specific primers. *Gapdh* amplification was used to normalize the samples.

### Actin polymerization assay

Repolymerization of the actin, depolymerized by oxidation, was examined with modification as described previously^39^. Briefly, pyrene-labelled actin was oxidized by treatment with hydrogen peroxide and then prepared for the polymerization assay by transferring it to polymerization buffer (Cytoskeleton, Inc). This oxidized pyrene-labelled actin, with or without MSRB3^WT^ or MSRB3^p.Cys89Gly^ (MSRB3^Mut^), was monitored for changes in fluorescence intensity to check polymerization in the presence of dithiothreitol [DTT]).

## Acknowledgements

We are very thankful to Drs. R. Yousaf, A.P.J. Giese, Y. Sokolova, and Ms. L.M. Schroeder-Carter for their technical assistance, and to Dr. D. Winkler, Director, Advanced Imaging Core, NIDCD for allowing us to use the JOEL2100 TEM microscope. This work was supported by R01DC011803 (SR), R01AG064223 (GV) and funded (in part) by the NIDCD DIR DC000039 to T.B.F.

## Author information

### Contributions

Conceived and designed the experiments: G.N., E.M.R., and B.C.L. Performed the experiments: G.N., E.M.R., B.C.L., G.P.R., I.A.B. and B.M. Analyzed the data: G.N., E.M.R., B.C.L. Contributed reagents and materials: T.B.F., V.N.G. and S.R. Discussed the results and commented on the manuscript: G.N., E.M.R., T.B.F., V.N.G. and S.R. Drafted and revised the paper: G.N., E.M.R., and S.R.

### Data availability

The data and analysis are available upon reasonable request from the corresponding author (S.R.).

## References

1. Huang, T., Cheng, A.G., Stupak, H., Liu, W., Kim, A., Staecker, H., Lefebvre, P.P., Malgrange, B., Kopke, R., Moonen, G., et al. (2000). Oxidative stress-induced apoptosis of cochlear sensory cells: otoprotective strategies. Int J Dev Neurosci 18, 259–270.

2. Ahmed, Z.M., Yousaf, R., Lee, B.C., Khan, S.N., Lee, S., Lee, K., Husnain, T., Rehman, A.U., Bonneux, S., Ansar, M., et al. (2011). Functional null mutations of MSRB3 encoding methionine sulfoxide reductase are associated with human deafness DFNB74. Am J Hum Genet 88, 19–29.

3. Shafique, S., Siddiqi, S., Schraders, M., Oostrik, J., Ayub, H., Bilal, A., Ajmal, M., Seco, C.Z., Strom, T.M., Mansoor, A., et al. (2014). Genetic spectrum of autosomal recessive non-syndromic hearing loss in Pakistani families. PLoS One 9, e100146.

4. Richard, E.M., Santos-Cortez, R.L.P., Faridi, R., Rehman, A.U., Lee, K., Shahzad, M., Acharya, A., Khan, A.A., Imtiaz, A., Chakchouk, I., et al. (2019). Global genetic insight contributed by consanguineous Pakistani families segregating hearing loss. Hum Mutat 40, 53–72.

5. Alqudah, S., Chertoff, M., Durham, D., Moskovitz, J., Staecker, H., and Peppi, M. (2018). Methionine Sulfoxide Reductase A Knockout Mice Show Progressive Hearing Loss and Sensitivity to Acoustic Trauma. Audiol Neurootol 23, 20–31.

6. Kwon, T.J., Cho, H.J., Kim, U.K., Lee, E., Oh, S.K., Bok, J., Bae, Y.C., Yi, J.K., Lee, J.W., Ryoo, Z.Y., et al. (2014). Methionine sulfoxide reductase B3 deficiency causes hearing loss due to stereocilia degeneration and apoptotic cell death in cochlear hair cells. Hum Mol Genet 23, 1591–1601.

7. Levine, R.L., Mosoni, L., Berlett, B.S., and Stadtman, E.R. (1996). Methionine residues as endogenous antioxidants in proteins. Proc Natl Acad Sci U S A 93, 15036–15040.

8. Keck, R.G. (1996). The use of t-butyl hydroperoxide as a probe for methionine oxidation in proteins. Anal Biochem 236, 56–62.

9. Brot, N., and Weissbach, H. (1983). Biochemistry and physiological role of methionine sulfoxide residues in proteins. Arch Biochem Biophys 223, 271–281.

10. Vogt, W. (1995). Oxidation of methionyl residues in proteins: tools, targets, and reversal. Free Radic Biol Med 18, 93–105.

11. Hoshi, T., and Heinemann, S. (2001). Regulation of cell function by methionine oxidation and reduction. J Physiol 531, 1–11.

12. Weissbach, H., Etienne, F., Hoshi, T., Heinemann, S.H., Lowther, W.T., Matthews, B., St John, G., Nathan, C., and Brot, N. (2002). Peptide methionine sulfoxide reductase: structure, mechanism of action, and biological function. Arch Biochem Biophys 397, 172–178.

13. Kim, H.Y., and Gladyshev, V.N. (2004). Methionine sulfoxide reduction in mammals: characterization of methionine-R-sulfoxide reductases. Mol Biol Cell 15, 1055–1064.

14. Sharov, V.S., Ferrington, D.A., Squier, T.C., and Schoneich, C. (1999). Diastereoselective reduction of protein-bound methionine sulfoxide by methionine sulfoxide reductase. FEBS Lett 455, 247–250.

15. Moskovitz, J., Poston, J.M., Berlett, B.S., Nosworthy, N.J., Szczepanowski, R., and Stadtman, E.R. (2000). Identification and characterization of a putative active site for peptide methionine sulfoxide reductase (MsrA) and its substrate stereospecificity. J Biol Chem 275, 14167–14172.

16. Moskovitz, J., Singh, V.K., Requena, J., Wilkinson, B.J., Jayaswal, R.K., and Stadtman, E.R. (2002). Purification and characterization of methionine sulfoxide reductases from mouse and Staphylococcus aureus and their substrate stereospecificity. Biochem Biophys Res Commun 290, 62–65.

17. Etienne, F., Spector, D., Brot, N., and Weissbach, H. (2003). A methionine sulfoxide reductase in Escherichia coli that reduces the R enantiomer of methionine sulfoxide. Biochem Biophys Res Commun 300, 378–382.

18. Vougier, S., Mary, J., and Friguet, B. (2003). Subcellular localization of methionine sulphoxide reductase A (MsrA): evidence for mitochondrial and cytosolic isoforms in rat liver cells. Biochem J 373, 531–537.

19. Kim, H.Y., and Gladyshev, V.N. (2004). Characterization of mouse endoplasmic reticulum methionine-R-sulfoxide reductase. Biochem Biophys Res Commun 320, 1277–1283.

20. Shen, X., Liu, F., Wang, Y., Wang, H., Ma, J., Xia, W., Zhang, J., Jiang, N., Sun, S., Wang, X., et al. (2015). Down-regulation of msrb3 and destruction of normal auditory system development through hair cell apoptosis in zebrafish. Int J Dev Biol 59, 195–203.

21. Kim, M.A., Cho, H.J., Bae, S.H., Lee, B., Oh, S.K., Kwon, T.J., Ryoo, Z.Y., Kim, H.Y., Cho, J.H., Kim, U.K., et al. (2016). Methionine Sulfoxide Reductase B3-Targeted In Utero Gene Therapy Rescues Hearing Function in a Mouse Model of Congenital Sensorineural Hearing Loss. Antioxid Redox Signal 24, 590–602.

22. Yancopoulos, G.D. (2003). VelociGene: a high-throughput approach for functionizing the genome via custom gene mutation and high-resolution expression analysis in mice. Breast Cancer Research 5, 56.

23. Pacentine, I., Chatterjee, P., and Barr-Gillespie, P.G. (2020). Stereocilia Rootlets: Actin-Based Structures That Are Essential for Structural Stability of the Hair Bundle. Int J Mol Sci 21.

24. Kitajiri, S., Fukumoto, K., Hata, M., Sasaki, H., Katsuno, T., Nakagawa, T., Ito, J., Tsukita, S., and Tsukita, S. (2004). Radixin deficiency causes deafness associated with progressive degeneration of cochlear stereocilia. J Cell Biol 166, 559–570.

25. Lee, B.C., Peterfi, Z., Hoffmann, F.W., Moore, R.E., Kaya, A., Avanesov, A., Tarrago, L., Zhou, Y., Weerapana, E., Fomenko, D.E., et al. (2013). MsrB1 and MICALs regulate actin assembly and macrophage function via reversible stereoselective methionine oxidation. Mol Cell 51, 397–404.

26. Kim, H.Y., and Gladyshev, V.N. (2007). Methionine sulfoxide reductases: selenoprotein forms and roles in antioxidant protein repair in mammals. Biochem J 407, 321–329.

27. Lee, B.C., Dikiy, A., Kim, H.Y., and Gladyshev, V.N. (2009). Functions and evolution of selenoprotein methionine sulfoxide reductases. Biochim Biophys Acta 1790, 1471–1477.

28. Katsuno, T., Belyantseva, I.A., Cartagena-Rivera, A.X., Ohta, K., Crump, S.M., Petralia, R.S., Ono, K., Tona, R., Imtiaz, A., Rehman, A., et al. (2019). TRIOBP-5 sculpts stereocilia rootlets and stiffens supporting cells enabling hearing. JCI Insight 4.

29. Krey, J.F., Liu, C., Belyantseva, I.A., Bateschell, M., Dumont, R.A., Goldsmith, J., Chatterjee, P., Morrill, R.S., Fedorov, L.M., Foster, S., et al. (2022). ANKRD24 organizes TRIOBP to reinforce stereocilia insertion points. J Cell Biol 221.

30. Drummond, M.C., Belyantseva, I.A., Friderici, K.H., and Friedman, T.B. (2012). Actin in hair cells and hearing loss. Hear Res 288, 89–99.

31. Tilney, L.G., Egelman, E.H., DeRosier, D.J., and Saunder, J.C. (1983). Actin filaments, stereocilia, and hair cells of the bird cochlea. II. Packing of actin filaments in the stereocilia and in the cuticular plate and what happens to the organization when the stereocilia are bent. J Cell Biol 96, 822–834.

32. Tilney, L.G., and DeRosier, D.J. (1986). Actin filaments, stereocilia, and hair cells of the bird cochlea. IV. How the actin filaments become organized in developing stereocilia and in the cuticular plate. Dev Biol 116, 119–129.

33. Kitajiri, S., Sakamoto, T., Belyantseva, I.A., Goodyear, R.J., Stepanyan, R., Fujiwara, I., Bird, J.E., Riazuddin, S., Riazuddin, S., Ahmed, Z.M., et al. (2010). Actin-bundling protein TRIOBP forms resilient rootlets of hair cell stereocilia essential for hearing. Cell 141, 786–798.

34. Manor, U., and Kachar, B. (2008). Dynamic length regulation of sensory stereocilia. Semin Cell Dev Biol 19, 502–510.

35. Zhang, D.S., Piazza, V., Perrin, B.J., Rzadzinska, A.K., Poczatek, J.C., Wang, M., Prosser, H.M., Ervasti, J.M., Corey, D.P., and Lechene, C.P. (2012). Multi-isotope imaging mass spectrometry reveals slow protein turnover in hair-cell stereocilia. Nature 481, 520–524.

36. Narayanan, P., Chatterton, P., Ikeda, A., Ikeda, S., Corey, D.P., Ervasti, J.M., and Perrin, B.J. (2015). Length regulation of mechanosensitive stereocilia depends on very slow actin dynamics and filament-severing proteins. Nat Commun 6, 6855.

37. Drummond, M.C., Barzik, M., Bird, J.E., Zhang, D.S., Lechene, C.P., Corey, D.P., Cunningham, L.L., and Friedman, T.B. (2015). Live-cell imaging of actin dynamics reveals mechanisms of stereocilia length regulation in the inner ear. Nat Commun 6, 6873.

38. Hung, R.J., Yazdani, U., Yoon, J., Wu, H., Yang, T., Gupta, N., Huang, Z., van Berkel, W.J., and Terman, J.R. (2010). Mical links semaphorins to F-actin disassembly. Nature 463, 823–827.

39. Hung, R.J., Pak, C.W., and Terman, J.R. (2011). Direct redox regulation of F-actin assembly and disassembly by Mical. Science 334, 1710–1713.

40. Giridharan, S.S., Rohn, J.L., Naslavsky, N., and Caplan, S. (2012). Differential regulation of actin microfilaments by human MICAL proteins. J Cell Sci 125, 614–624.

41. Moskovitz, J., Bar-Noy, S., Williams, W.M., Requena, J., Berlett, B.S., and Stadtman, E.R. (2001). Methionine sulfoxide reductase (MsrA) is a regulator of antioxidant defense and lifespan in mammals. Proc Natl Acad Sci U S A 98, 12920–12925.

42. Fomenko, D.E., Novoselov, S.V., Natarajan, S.K., Lee, B.C., Koc, A., Carlson, B.A., Lee, T.H., Kim, H.Y., Hatfield, D.L., and Gladyshev, V.N. (2009). MsrB1 (methionine-R-sulfoxide reductase 1) knock-out mice: roles of MsrB1 in redox regulation and identification of a novel selenoprotein form. J Biol Chem 284, 5986–5993.

43. Martinez, Y., Li, X., Liu, G., Bin, P., Yan, W., Mas, D., Valdivie, M., Hu, C.A., Ren, W., and Yin, Y. (2017). The role of methionine on metabolism, oxidative stress, and diseases. Amino Acids 49, 2091–2098.

44. Giese, A.P.J., Tang, Y.Q., Sinha, G.P., Bowl, M.R., Goldring, A.C., Parker, A., Freeman, M.J., Brown, S.D.M., Riazuddin, S., Fettiplace, R., et al. (2017). CIB2 interacts with TMC1 and TMC2 and is essential for mechanotransduction in auditory hair cells. Nat Commun 8, 43.

