## Supplementary Figures for "Methionine sulfoxide reductase B3 antioxidant activity is indispensable for mouse inner ear cuticular plate structure and hair bundle integrity"

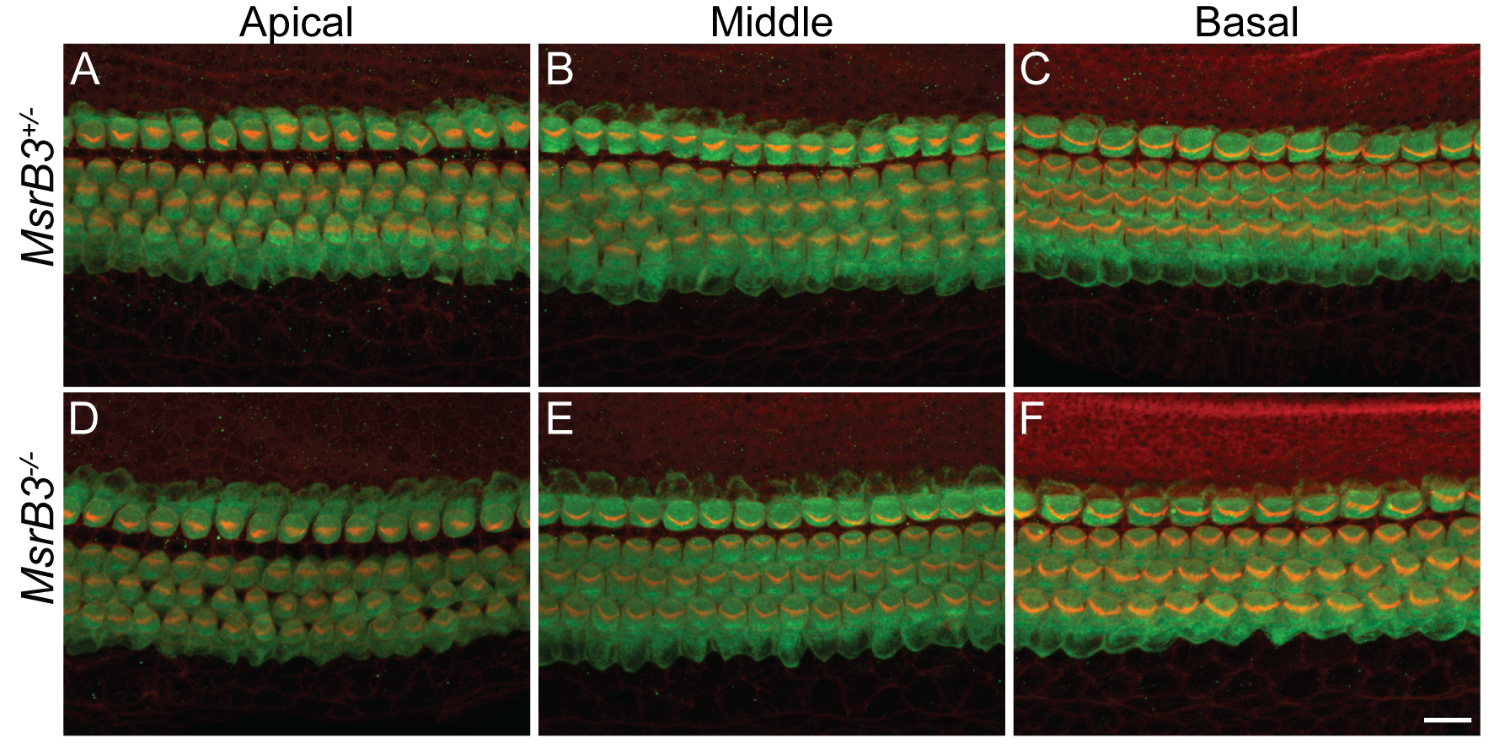

**Supplementary figure 1. Hair cells of *MsrB3^-/-^* mutant mice appear to have normal development and morphology at P0.** Maximum intensity projections of confocal Z-stacks from the apical, middle and basal turns of the organ of Corti whole-mount cochleae labeled with anti-β-spectrin (green) and phalloidin (red) are shown for normal hearing heterozygous (A-C) and *MsrB3* mutant mice (D-F) at P0. Scale bar is 5µm and applies to all panels.

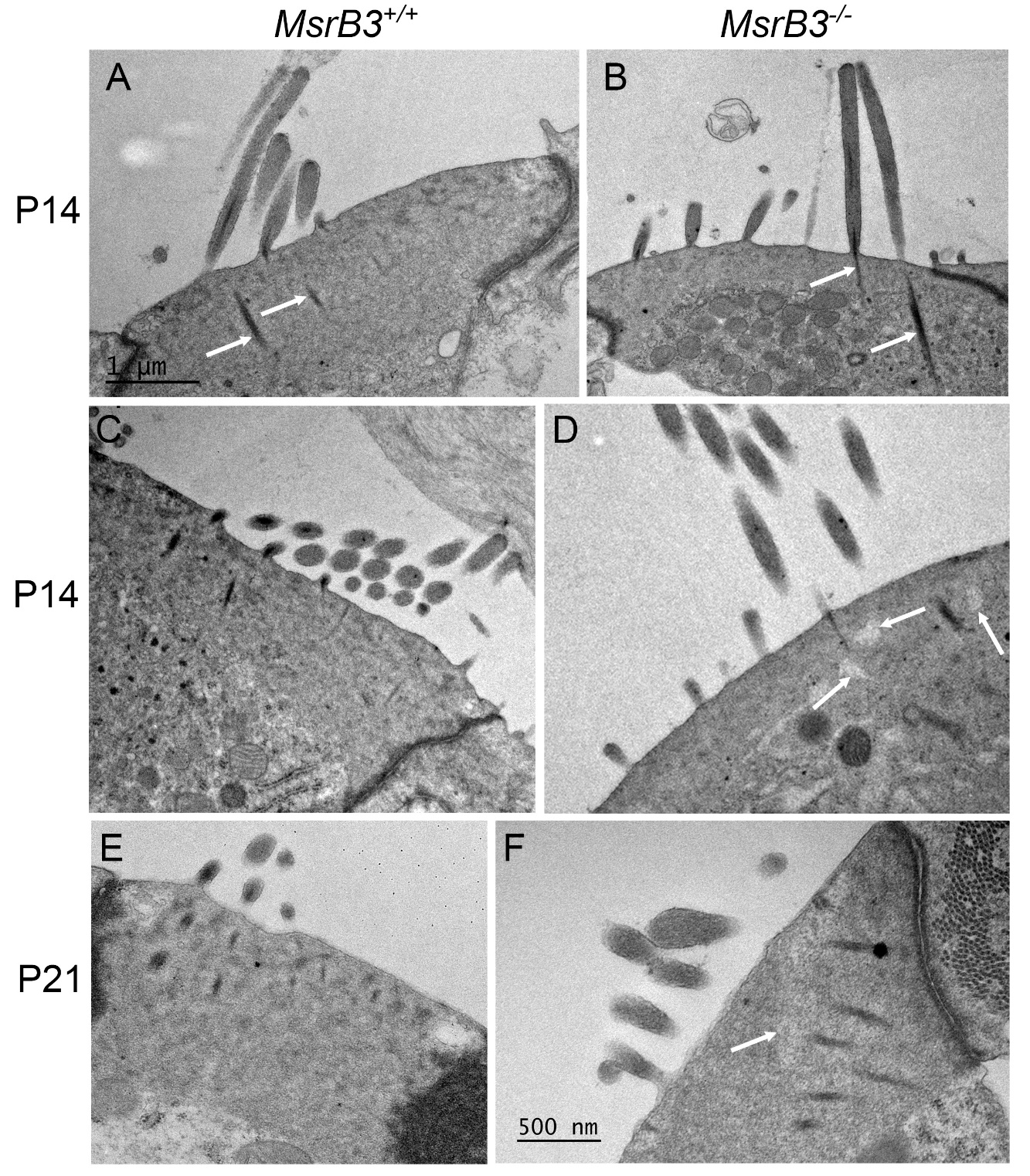

**Supplementary figure 2. *MsrB3^-/-^* mutant mice did not exhibit obvious stereocilia rootlet alterations but show uneven density of the cuticular plate.** Transmission electron micrographs of control (A, C) and *MsrB3* mutant mouse OHCs (B, D) at P14. Arrows in A and B point to stereocilia rootlets present in both wild-type and mutant hair cells. (C) Control P14 wild-type OHC cuticular plate shows relatively homogeneous density around more electron dense rootlets, while P14 mutant OHC cuticular plate (D) shows uneven density in rootlet proximities with lighter areas indicated by arrows. Similar changes in density of the cuticular plate were observed in P21 mutant OHCs (F, arrow) as compared to P21 wild-type OHC (E). Scale bars: 1µm applies to A-E, and 500 nm to F.

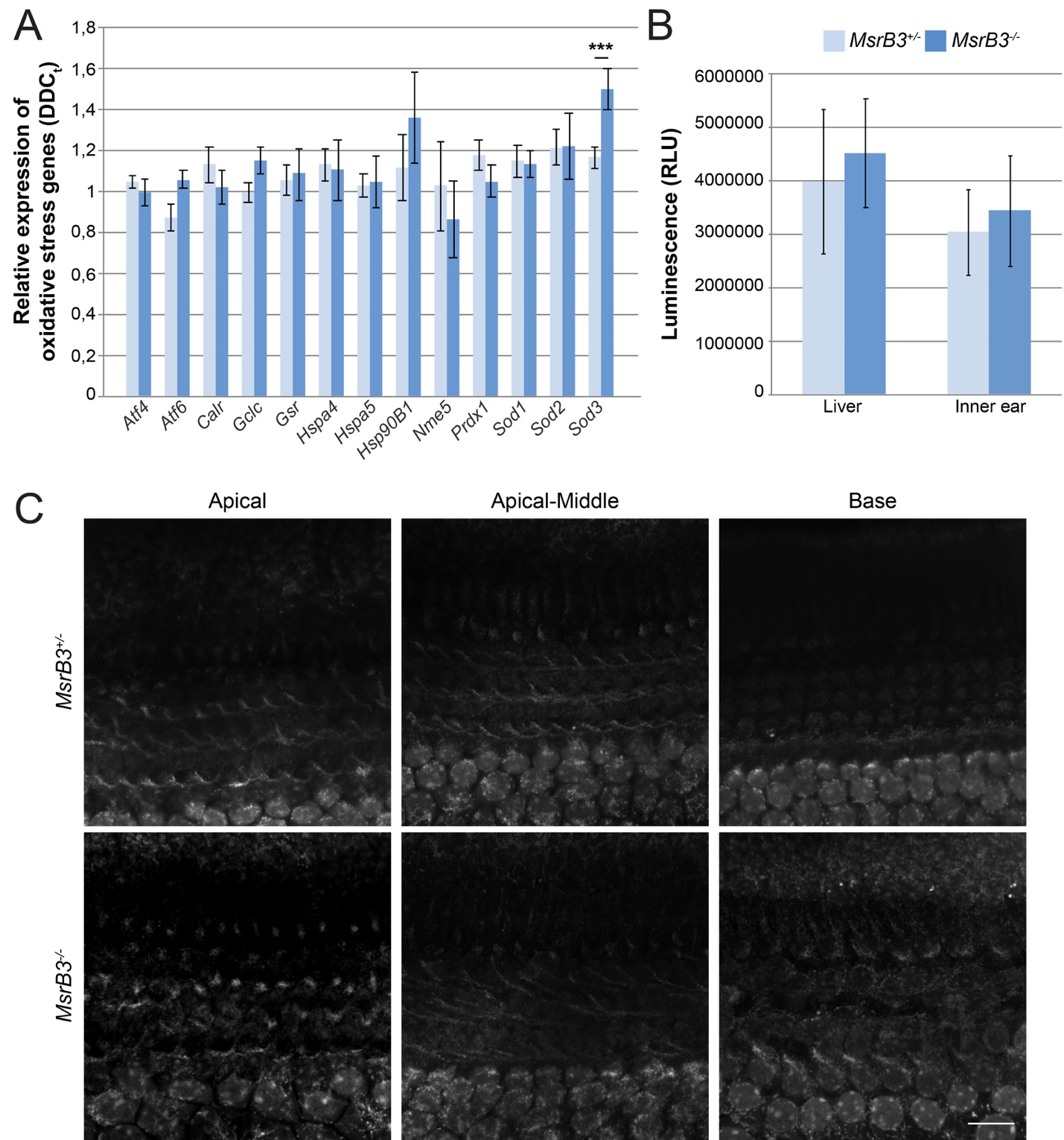

**Supplementary figure 3. Characterization of oxidative stress in MsrB3 mice inner ear.** (A) Quantification of oxidative stress representative genes expression, normalized against *Gapdh*, in the cochleae of *MsrB3^-/-^* mutant mice and their control littermates (*MsrB3^+/-^*) at P15. No variation of expression was observed except for *SOD3* (means ± SEM). ∗∗∗P<0.001; unpaired t-test. (B) Quantification of total glutathione in relative luminescence units (RLU) from mutant *MsrB3^-/-^* and control *MsrB3^+/-^* liver and inner ear protein extracts at P14 (means ± SEM). (C) Representative confocal images of CellROX fluorescence displaying production of ROS in mutant *MsrB3^-/-^* and control *MsrB3^+/-^* inner ear explant. No differences were observed between mutant and control conditions. Scale bar: 15µm.
